# Saddle Buckling of a Multicellular Alga Driven by Differential Growth

**DOI:** 10.64898/2026.11.20.767507

**Authors:** Zhao Wang, Sumit Kumar Birwa, Jane Y. Y. Chui, Kyriacos C. Leptos, Raymond E. Goldstein

## Abstract

During development of many simple multicellular organisms there is a complex interplay between cell growth and generation of an extracellular matrix that confers structure. We study this problem quantitatively using *Gonium pectorale*, a 16-cell alga that transforms from a flat to a buckled shape during growth without cell division or rearrangement. Cell-level measurements and 3D reconstructions show that the 12 peripheral cells grow faster than the central 4, producing hoop stress that selects the saddle mode, whose amplitude displays a continuous bifurcation with classical square root singularity. These results are consistent with a continuum model of differential growth within Föppl–von Kármán theory, which yields information on the mechanical properties of the ECM. Implications of this transformation in evolutionary transitions to multicellularity are discussed.

---

In the long history of scientific studies of organism growth, it was J.S. Huxley [1] who perhaps first clearly articulated the existence of “differential growth” as a key feature. Inspired by the earlier geometric analyses by D’Arcy Thompson [2], across a wide range of animals Huxley provided empirical evidence not only for organs growing at different rates within an organism (e.g., the claw of the fiddler crab relative to the rest of its body), but also for “growth gradients” within a given organ (e.g., between the proximal and distal regions of the claw).

Only much more recently have the mechanical implications of differential growth on the three-dimensional shapes of continua been understood. Within elasticity theory, incompatible local growth rates create residual stress; for ostensibly flat, sheet-like tissues, that stress can be relieved by out-of-plane bending [3–7], as in patterned gels [8–10] and in growing leaves, flowers, and the gut [11–16]. The response of the tissue depends on the spatial distribution of the growth and on the nature of the mechanical properties of the tissue.

These issues take on particular significance when considering simple organisms that serve as models for evolutionary transitions to multicellularity. Both for Volvocine green algae (spanning from unicellular *Chlamydomonas* to *Volvox*) and choanoflagellates (the closest living relatives of animals), cells are typically held together by an extracellular matrix (ECM). As this provides the physical scaffold that dictates overall morphology, and directly enables specialized behaviours like swimming, feeding, and phototaxis, proper orchestration of cell growth and ECM generation is crucial. In higher organisms, tissue growth is often accompanied by cell division, migration, and rearrangement, making these two contributions difficult to separate [17]; it is thus of general interest to identify a fixed cellular assembly amenable to precise measurements.

Here we show that the 16-cell volvocine alga *Gonium pectorale* [18–20], whose life cycle is shown in Fig. 1, provides such an assembly. Once hatched, the juvenile is a flat plate whose 4 central and 12 peripheral cells (“4 + 12”) grow while held together with an ECM, eventually buckling. Later, each cell divides to produce 16 new 16-cell progeny, with 12 in a pinwheel arrangement around the central 4 (Fig. 1, left). This cycle provides a minimal setting to examine how differential cell growth is transmitted through a mechanically connected ECM: its small, fixed architecture permits cell-resolved growth measurements, while micropipette aspiration allows continuous, long-term observation of colony shape.

**FIG. 1.**
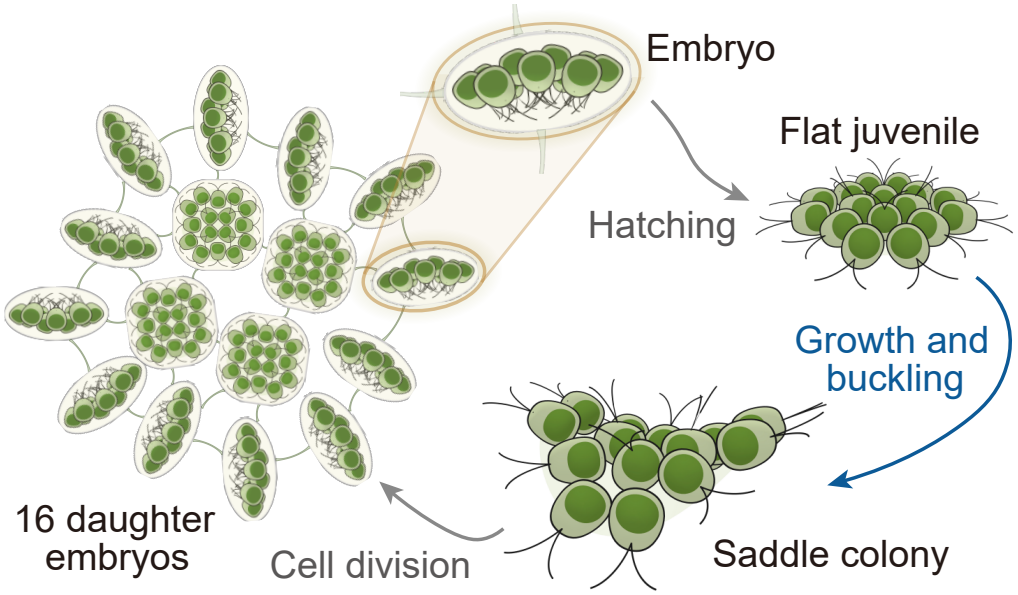
Life cycle of *G. pectorale*. Each cell divides into a 16-cell daughter embryos which hatch as flat juveniles that grow and buckle into a saddle (this work) before dividing again.

Although “warping” of *Gonium* colonies was noted long ago [21], this feature has not been quantified or explained. We find that peripheral cells grow faster than central ones, and show that a two-region elastic model [Fig. 3(a)] converts this imbalance into hoop compression that selects the saddle mode independently, identified by 3D reconstruction. The continuous onset with exponent 0.52 *±* 0.05 is consistent with the predicted bifurcation. With the measured critical size as input, the growth records account for the near-onset amplitude without fitting. The growth pattern thus selects the saddle, while the mechanics of the ECM-connected assembly sets how much growth is needed for it to appear: the onset serves as a mechanical assay, rationalizing an effective mechanical thickness associated with the ECM.

## Shape transition

Wild-type colonies of *G. pectorale* (CCAC strain 3275 B) were grown axenically in standard *Volvox* medium [22] at 22 ^°^C under ∼3800 lx illumination on a 14 h*/*10 h light/dark cycle. Each newly formed 16-cell colony was held via one peripheral cell by gentle aspiration onto a polished glass micropipette of ∼3 *µ*m inner diameter, mounted on a rotation stage to enable both face-on and edge-on imaging. The transition occurs during early growth, while the ECM appears continuous and the cell number and 4 + 12 arrangement are fixed (See § IA of Supplemental Material [23] for details).

Figure 2(a) shows an initially flat colony developing a large out-of-plane deformation over 29 h (Movie 1 in [23]). We enclose the colony body in each image by its minimum-area bounding rectangle, with short and long sides *b* and *L* and aspect ratio *a* = *b/L*, and let

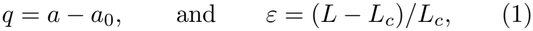

where *a*_0_ is the pre-transition plateau of *a* and the critical size *L*_*c*_ (§ IB in [23]) is that at which buckling first appears. Across 6 colonies, 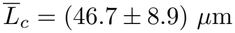 (where all uncertainties herein are standard deviations). The amplitude *q* rises from zero continuously, with no detectable jump at onset. Fitting *q* ∝ *ε*^*β*^ to each colony gives *β* = 0.52 *±* 0.05. A separate square-root fit gives a prefactor *C* for each colony. The normalized amplitude *Q* = *q/C* collapses the data onto 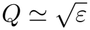 [Fig. 2(b)]. The continuous square-root growth of *q* is consistent with a supercritical pitchfork bifurcation, confirmed below by the sign of the cubic term in an amplitude equation.

**FIG. 2.**
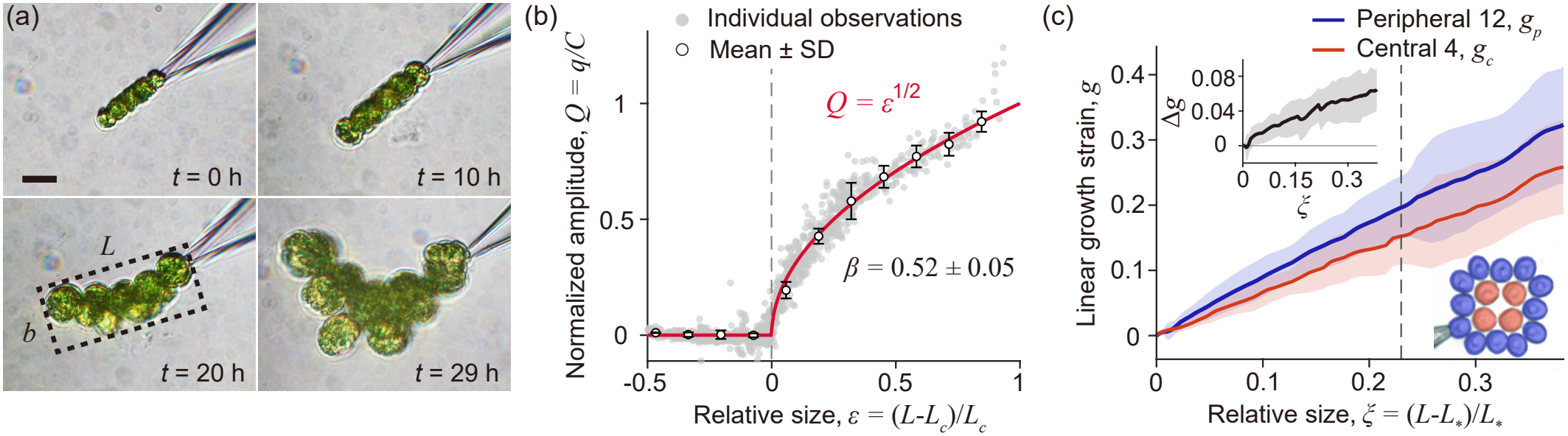
Flat-to-saddle transition of growing *G. pectorale*. (a) Edge-on time-lapse of a colony held by a micropipette, with dashed minimum-area bounding rectangle in *t* = 20 h frame. Scale bar: 10 *µ*m. (b) Normalized shape amplitude versus relative size for six colonies, where *C* is a colony-specific constant; *β* is obtained from per-colony fits of *q ∝ ϵ*^*β*^. (c) Cumulative linear growth strain of peripheral (blue) and central (red) cells from 6 face-on growth records. The dashed line marks the mean critical size 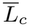 of the shape cohort. Upper inset: mismatch Δ*g* = *g*_*p*_ *− g*_*c*_. Lower inset: central and peripheral cell groups.

## Differential growth

A separate set of 6 colonies was imaged face-on in bright-field *z* stacks, with each cell’s projected area measured in its best-focused plane (§ IC in [23]). For each colony, let *S*_*α*_ (for *α* = *c, p*) denote the mean area of the central 4 or peripheral 12 cells [Fig. 2(c)]. We define the regional linear expansion and relative size as

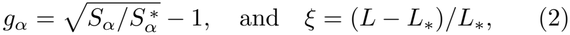

where 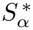 is the reference mean area at *L*_*_, the smallest colony size covered by all six records. The square root converts the area ratio into a length ratio, as the growth tensor in the model describes expansion of line elements.

Figure 2(c) shows greater relative expansion of peripheral cells and an overall increase in the mismatch Δ*g* = *g*_*p*_ − *g*_*c*_. At the mean critical size 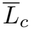 obtained from the shape measurements (dashed line), the differential growth is Δ*g* = 0.065 *±* 0.028. Growth thus builds an increasing mismatch across the fixed 4 + 12 arrangement, providing a measured input for the elastic analysis.

## Theory

We model the colony as a thin 2-region elastic disk of radius *R* = *L/*2 and thickness *H*, with radial coordinate *r* and internal division at *r* = *λR*. Setting *λ* = 1*/*2 assigns one quarter of the disk area to the center [Fig. 3(a)], as in the 4:12 partition. Each region grows locally isotropically, by *g*_c_ or *g*_p_. In the mechanical analysis, strains are referenced to each colony’s first recorded planar state *L*_0_, with no additional initial mismatch. The sheet has uniform 2D Young modulus *K* (*K* = *HY*, where *Y* is the 3D Young’s modulus), bending rigidity *D*, Poisson ratio *ν* = 1*/*2 and free outer boundary.

**FIG. 3.**
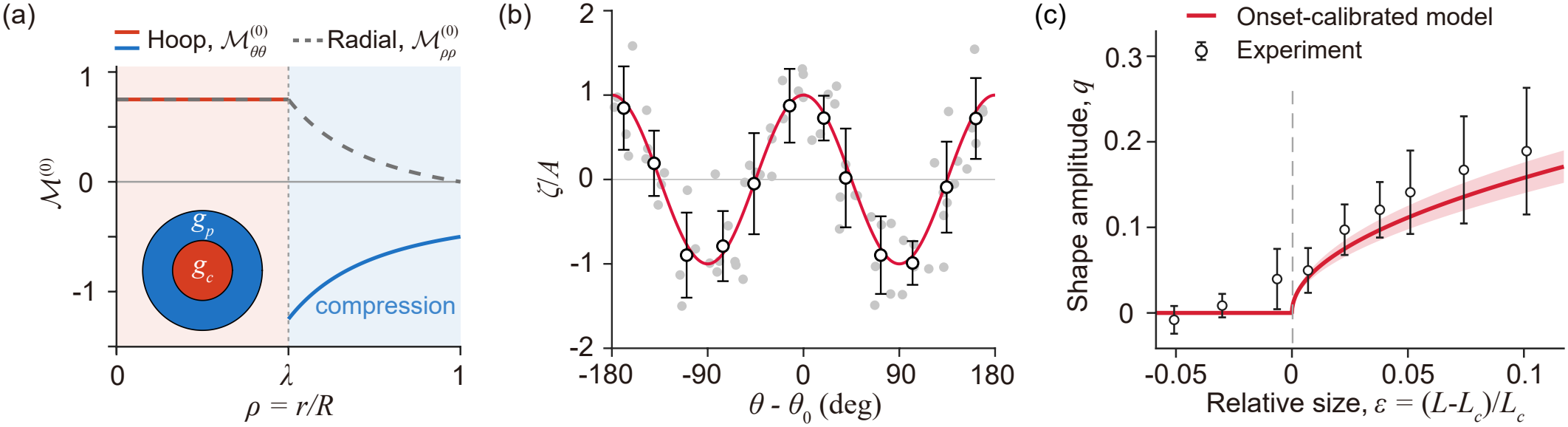
Growth-induced stress, saddle symmetry, and near-onset amplitude. (a) Base-state stress resultants *M*^(0)^ of the two-region disk for *λ* = 1*/*2: radial (dashed) and hoop (solid). Inset: growth regions. (b) Peripheral cell heights of 6 colonies relative to their best-fit planes, each rotated by its own *m* = 2 phase *θ*_0_ and normalized by its fitted amplitude *A* (grey). Circles show 30^*°*^ bins; red line is the mode shape cos 2(*θ − θ*_0_). (c) Amplitude predicted from the six measured growth curves and independently measured side-view amplitudes, shown over their common near-onset range.

Differential growth between the central and peripheral regions leads to longitudinal stresses that result in out-of-plane displacements *ζ*(*r, θ*). Anticipating a perturbative expansion of the stress tensor *N* in powers of *ζ*, we consider first the longitudinal stresses 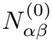 in a flat disk in the membrane limit [24], where the 2D elastic strain tensor *E* is related to the displacement **u** and growth rate *g* via *E*_*αβ*_ = (*u*_*α,β*_ + *u*_*β,α*_)*/*2 − *gδ*_*αβ*_. With azimuthal symmetry and radial displacement 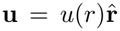, the stress resultants 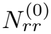 and 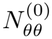 are given by Eq. (8) in End Matter, with 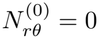. The equilibrium condition 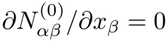 reduces to *u*^*′′*^ +*u*^*′*^*/r* − *u/r*^2^ = 0, with regularity at *r* = 0, a free edge at *r* = *R*, and continuity in *u* and 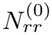 at *r* = *λR*. With the rescalings 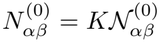 and *ρ* = *r/R* we find 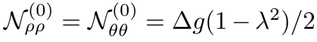 in the center and in the periphery (*λ < ρ* ≤ 1)

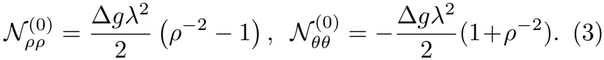

As all the nonzero components of *N* ^(0)^ are proportional to Δ*g/*2, we let 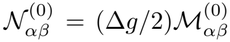. For Δ*g >* 0 the center is under uniform biaxial tension, and although growth is isotropic, 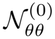 jumps at *ρ* = *λ* and is compressive throughout the outer annulus [Fig. 3(a)]; as the two regions are connected, the faster-growing periphery cannot expand freely in the circumferential direction.

These results derive from an Airy stress function *χ* via 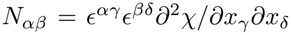, and scaled Φ = *χ/KR*^2^. In the center, we find Φ^(0)^ = Δ*g*(1 −*λ*^2^)*ρ*^2^*/*4, while in the periphery Φ^(0)^ = (Δ*gλ*^2^*/*2)(ln(*ρ/λ*) − *ρ*^2^*/*2 + 1*/*2). Now allowing for non-zero *ζ*, we set 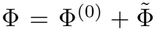, where 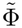 is the correction. The spatially nonuniform growth is accounted for by the flat-state stress function Φ^(0)^. Subtracting this, the growth-modified Föppl–von Kármán equations [6] are coupled PDEs for the scaled displacement *w* = *ζ/R* and 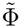. Defining Γ = *D/*(*KR*^2^), they are

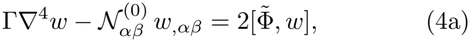

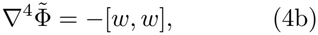

where [*f, g*] = (*f*_*xx*_*g*_*yy*_ + *f*_*yy*_*g*_*xx*_ − 2*f*_*xy*_*g*_*xy*_)*/*2, and [*w, w*] is the Gaussian curvature of the sheet (§ IIA in [23]).

Linearizing Eq. (4) about the flat state removes the right-hand side of its first line, which is cubic in *w* because 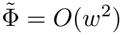. The linear problem becomes

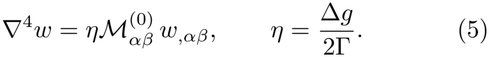

Eq. (5) depends on growth and mechanics only through *η*; it must be solved in each region with boundary conditions at *ρ* = 0, *λ*, 1, and *η* = *η*_*c*_(*m*) is an eigenvalue for modes *w* ∝ *ϕ*_*m*_(*ρ*) cos(*mθ*). For *λ* = *ν* = 1*/*2, the first few numerically obtained thresholds [23] are *η*_*c*_(2) = 9.16 (a saddle), *η*_*c*_(3) = 19.4, *η*_*c*_(4) = 33.7. Axisymmetric buckling occurs if the center outgrows the periphery (Δ*g <* 0).

In our experiments, we identify the mode from cell-center positions reconstructed from the face-on *z* stacks (§ ID and Movie 2 in [23]). The displacements of peripheral cells from a best-fit plane, aligned in phase and normalized by amplitude (*ζ/A*), follow the azimuthal variation of the predicted saddle [Fig. 3(b)]. The significance of this fit is quantified by *P*_*m*_, the fraction of the squared peripheral height signal explained by a separate least-squares fit of mode *m*. Across the 6 colonies, *m* = 2 is the largest mode in every colony and *P*_2_ = (73 *±* 9)%. The observed spatial symmetry therefore matches the first instability generated by the measured growth pattern.

## Onset and amplitude

Buckling sets in when *η* = *η*_*c*_(2), so the measured critical mismatch Δ*g*_*c*_ determines the onset value of the stiffness ratio via Γ_*c*_ = Δ*g*_*c*_*/*2*η*_*c*_(2). This makes buckling a mechanical assay for Γ. Because growth and shape were recorded in separate cohorts, we evaluate each growth curve at the mean critical size 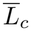 of the shape cohort. The six resulting estimates give 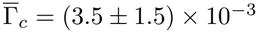, which we discuss below.

Beyond onset, the amplitude is limited by the nonlinearities in Eq. (4). Near onset, *w* = *A ϕ*_*c*_(*ρ*) cos 2*θ* + *· · ·*, where *ϕ*_*c*_ ≡ *ϕ*_2_ is the critical mode, normalized so that max |*ϕ*_*c*_| = 1, and *A* is the dimensionless maximum rim deflection. Projecting Eq. (4) onto this mode gives

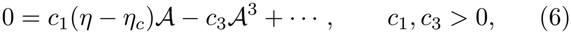

which contains only odd powers because (4) is invariant under *ζ* →−*ζ*. The cubic coefficient is positive because a saddle has nonzero Gaussian curvature and therefore cannot form without stretching the sheet, at an energetic cost of order *A*^4^ (End Matter). The amplitude therefore undergoes a supercritical pitchfork bifurcation, consistent with the continuous square-root onset in Fig. 2(b).

Converting *A* to the side-view observable gives *q*^2^ ≃ *C*_*g*_(Δ*g* −2*η*_*c*_Γ) above threshold, where *C*_*g*_ = *c*_1_*/*(2Γ*c*_3_) ≃ 1.08 is calculated from the mode shape and the observation geometry (§IIC in [23]). Over the near-onset window we take Γ ≃ Γ_*c*_, so that 2*η*_*c*_Γ ≃ Δ*g*_*c*_ and, for Δ*g >* Δ*g*_*c*_,

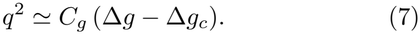

The growth records fix Δ*g* − Δ*g*_*c*_ and *C*_*g*_ is known, so Eq. (7) has no adjustable parameters. Since the measured mismatch increases approximately linearly with size near onset, Eq. (7) also predicts *q* ∝ *ε*^1*/*2^, consistent with the square-root onset observed in Fig 2(b).

We evaluate Eq. (7) for each growth record, with Δ*g*_*c*_ taken at 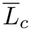, and average the amplitudes. Figure 3(c) compares these predictions with the experimental amplitudes over the common interval, −0.06 ≲ *ε* ≲ 0.12, with each shape record keeping its critical size. The prediction reproduces both the observed amplitude scale and its increase with *ε* within the between-colony spread.

With this effective description supported by the amplitude comparison, we compare the inferred stiffnesses with a homogeneous-sheet benchmark. For an incompressible homogeneous sheet whose thickness equals the cell-body diameter *H, D/K* = *H*^2^*/*9, which is a purely geometric result. With 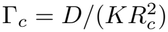 and 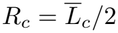, the colony-averaged *D/K* is smaller than this benchmark by a factor of ≃5.4, implying substantially weaker resistance to bending, relative to stretching, than for a homogeneous sheet of cell-body thickness. Equivalently, the mechanical thickness 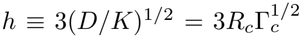, which equals *H* for a homogeneous sheet, is *h* = (0.42 *±* 0.10)*H*. Thus the buckling onset probes a load-bearing thickness less than half the colony thickness (the cell diameter *H*).

## Discussion

Thin intercellular ECM regions offer a possible explanation for this reduced ratio. For an elastic strip of thickness *h*, stretching stiffness scales ∼*h* and bending rigidity ∼*h*^3^, so thinning preferentially reduces bending resistance. We observe mesh-like ECM and elongated tethers after the saddle transition (§III in [23]). Thus, a nonuniform thickness may already be present in the ECM during buckling. A related distinction between geometric and effective bending thickness is incorporated in microtubule shell models to account for deformation in thin interprotofilament bridges [25]. If bending similarly localizes between *Gonium* cells, the effective mechanical thickness would depend on these compliant regions, even when cell bodies set the visible thickness. Resolving ECM thickness and deformation during the transition would directly test this interpretation.

Peripheral crowding offers a possible geometric link between saddle buckling and the subsequent colony development. Harper already noted [21] that the 12 peripheral cells occupy “the space which would be more naturally filled by ten”. Excess peripheral growth intensifies this crowding, and the sheet relieves it by bending out of plane. A similar out-of-plane accommodation reappears at division (Fig. 4): once each cell has formed a 16-cell daughter within the tethered matrix, the 12 peripheral daughters stand edge-on in a ring around the 4 central ones, which reorient only later. Together, these observations point to a common geometric principle: in an ECM-tethered colony, peripheral crowding is relieved out-of-plane, by collective bending of the sheet during growth and by reorientation of the connected daughters at division. This may be an adaptive trait that allows a multicellular organism to bear large multicellular progeny. A quantitative understanding of the late-stage geometry seen in Fig. 4 likely lies far outside the realm of Föpplvon Karman theory and would require a more intrinsic formulation of elastic sheet dynamics of the kind that has been described recently [26].

**FIG. 4.**
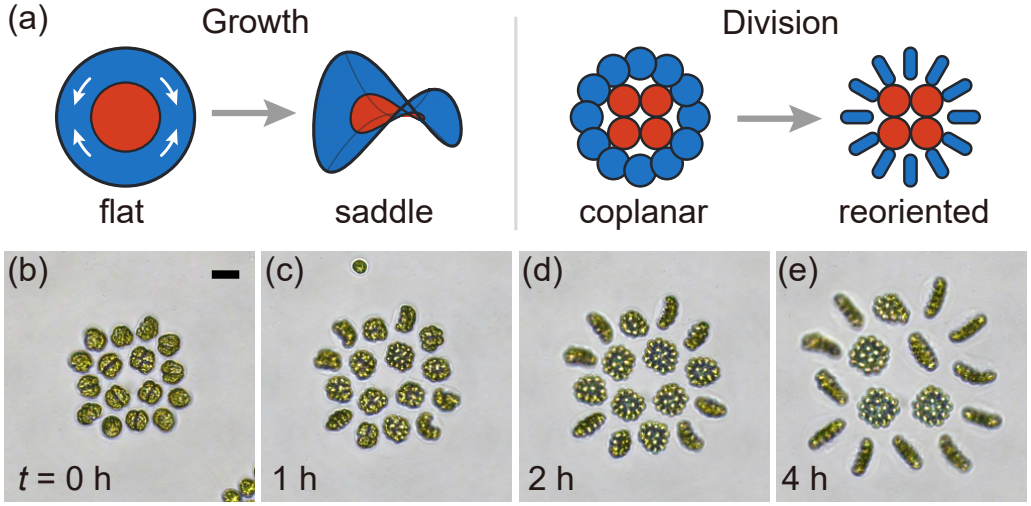
Peripheral crowding relieved out of plane. (a) Schematics of growth and division. (b)–(e) Time-lapse showing peripheral daughter colonies reorienting out of plane (§ IE in [23]). Scale bar, 10 *µ*m.

The comparison with uni- and multicellular relatives highlights a role of the ECM. In *Chlamydomonas*, the homologous material is a wall around each cell; the daughter cells formed within the mother-cell wall are released as independent individuals [27], so no shared matrix couples their growth. In *Volvox*, embryos develop inside glycoprotein vesicles whose controlled enlargement is required for inversion; when the vesicle fails to expand, inversion is impaired, and releasing the embryo restores it [28]. *Gonium* lies between these two: its cells, and later its daughters, are coupled through a common ECM, so growth of the parts generates stresses borne by the whole. The shape changes reported here are thus a direct mechanical consequence of multicellularity, of cells that grow while remaining attached. Tuning the growth pattern selects the mode, whereas remodelling of the matrix sets how much growth is needed for it to appear, offering two distinct routes by which an organism can control its form.

We thank Hester van Wachem for discussions. This work was supported in part by the Wellcome Trust Discovery Award 307079/Z/23/Z (ZW & JYYC), the Leverhulme Trust (KL & SKB) and the Complex Systems Fund at the University of Cambridge.

## Data Availability

The data that support the findings of this article are openly available [29].

## END MATTER

*Base state*—For a flat, axisymmetric colony, the displacement is radial, 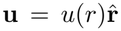, and the elastic strains are *E*_*rr*_ = *u*^*′*^ − *g* and *E*_*θθ*_ = *u/r* − *g*, where ^*′*^ = *d/dr* and *g* takes the constant value *g*_c_ or *g*_p_ in the respective regions. The 2D form of Hooke’s law for the sheet is

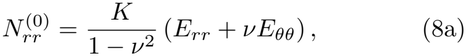

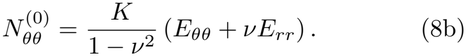

Radial force balance, 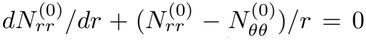, reduces in each region to *u*^*′′*^ + *u*^*′*^*/r* − *u/r*^2^ = 0. Thus, *u*(*r*) = *A*_*α*_*r* + *B*_*α*_*/r*, for *α* = c, p.

Regularity at the center requires *B*_c_ = 0, and the free outer edge requires 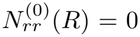, and at the bonded interface *r* = *λR* both *u* and, by radial force balance, 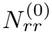 are continuous. These four conditions yield

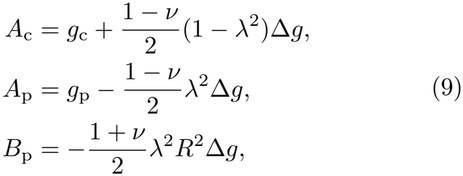

where Δ*g* = *g*_p_ −*g*_c_. Substituting these constants into Eqs. (8) and writing *ρ* = *r/R* and dividing by *K* gives the base stresses 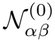 of the main text.

### Linear stability

For separable solutions to Eq. (5) of the form *w* = *ϕ*_*m*_(*ρ*) cos(*mθ*) we define *ℒ*_*m*_ = *d*^2^*/dρ*^2^ + *ρ*^−1^*d/dρ* − *m*^2^*/ρ*^2^ and use the result 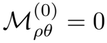 to obtain

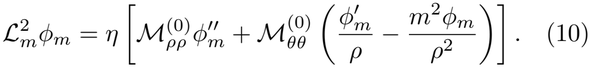

Substituting the components of *ℳ* ^(0)^ gives

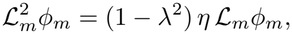

in the center, and in the periphery,

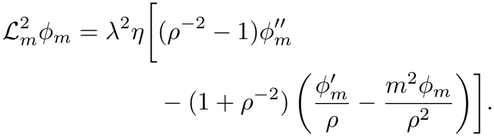

Growth and mechanics thus enter only through *η*.

Each regional equation is fourth order, giving four integration constants each in the center and the periphery. These eight conditions are: *(center)* The deflection *w* = *ϕ*_*m*_ cos(*mθ*) must be smooth and single-valued at the center. For *m* ≥ 2, this requires 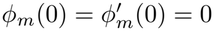; *(interface)* The sheet can neither tear nor fold along *ρ* = *λ*: its deflection and slope are continuous. No force or torque is applied along the interface, so the bending moment and transverse force are continuous. With *D* and *ν* equal in the two regions, and the radial tension continuous, these four conditions amount to continuity of *ϕ*_*m*_, 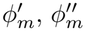 and 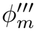 at *ρ* = *λ*; *(periphery)* This edge is free, so it carries no bending moment and no transverse force, the latter including the contribution of twisting along the edge. Substituting *w* = *ϕ*_*m*_ cos(*mθ*) into the standard expressions for these quantities [24], at *ρ* = 1, gives

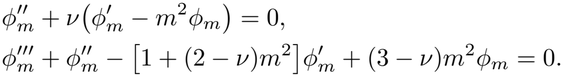

For each *m* ≥ 2, the eight conditions allow a nonzero deflection only at particular values of *η*. The smallest positive value defines *η*_*c*_(*m*). For *λ* = *ν* = 1*/*2, numerical solution (Chebyshev collocation) gives *η*_*c*_(2) = 9.16, *η*_*c*_(3) = 19.4 and *η*_*c*_(4) = 33.7, with the thresholds increasing monotonically up to *m* = 10. After removing rigid translation and tilt, no positive buckling eigenvalue was found for *m* = 0 or 1 (§ IIB in [23]). The *m* = 2 saddle is therefore the first unstable mode.

### Amplitude equation

Near onset we expand in the amplitude *A* of the critical mode *w*_1_ = *ϕ*_*c*_(*ρ*) cos 2*θ*, normalized by *ϕ*_*c*_(1) = 1. In an expansion in a formal small parameter *ϵ*, the leading deflection would be *ϵAw*_1_, so that *A* = *ϵA*. Equations (4) are invariant under *w* → − *w* with 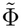 unchanged, so the deflection contains only odd powers of *A* and the induced stress function only even powers:

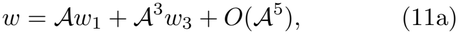

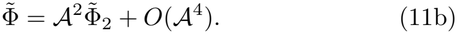

At fixed Γ the critical mismatch is Δ*g*_*c*_ = 2*η*_*c*_Γ. Since the base stress is proportional to Δ*g*, we have 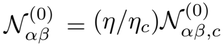, where *N* ^(0)^ is its value at onset. We take (*η* − *η*_*c*_)*/η*_*c*_ = *µA*^2^ + *O*(*A*^4^) and determine *µ* at order *A*^3^.

At *O*(*A*), Eq. (4a) gives the linear problem 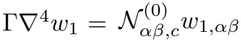, which determines the mode shape, leaving *A* unknown. At *O*(*A*^2^), Eq. (4b) gives

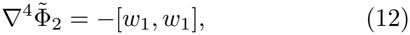

so the Gaussian curvature generates an induced stress function of *O*(*A*^2^). The function 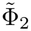 is smooth at the center and continuous with its first three radial derivatives at *ρ* = *λ*. At the traction-free outer edge *ρ* = 1, we choose the stress function to satisfy 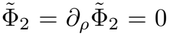. At *O*(*A*^3^), Eq. (4a) divided by Γ gives

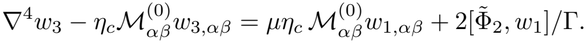

The operator on the left hand side is self-adjoint under the boundary and matching conditions and annihilates *w*_1_. Multiplying by *w*_1_ and integrating over the unit disk, with *dS* = *ρ dρ dθ*, eliminates *w*_3_: the right side must have zero projection onto *w*_1_. Integrating the prestress term by parts and using *η* − *η*_*c*_ ≃ *η*_*c*_*µA*^2^, we find

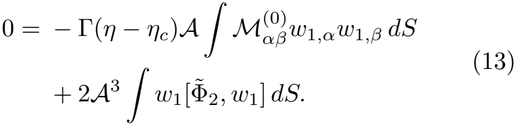

Comparing with Eq. (6), we obtain

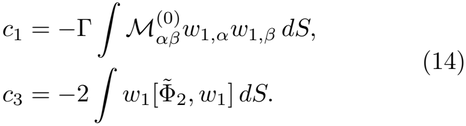

For *λ* = *ν* = 1*/*2, *c*_1_*/*2Γ ≃ 0.680 and *c*_3_ ≃ 0.630; since *q* ≃ |*A*| in the edge-on geometry [23], this gives *C*_*g*_ = *c*_1_*/*(2Γ*c*_3_) ≃ 1.08.

With the in-plane displacement relaxed, the elastic energy relative to the flat state is, in units of *KR*^2^,

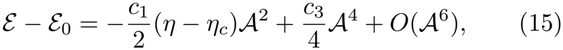

whose stationarity condition is Eq. (6). The quadratic term combines the bending cost with the energy released from the prestress, which balance at onset; the quartic term is the stretching cost of the saddle. The flat state is stable below onset and unstable above it, where two minima *A* = *±* [(*c*_1_*/c*_3_)(*η* −*η*_*c*_)]^1*/*2^, corresponding to opposite deflections, emerge continuously: a supercritical pitchfork bifurcation.

## SUPPLEMENTAL MATERIAL

### I. EXPERIMENTAL METHODS

#### A. Culture, micropipette holding, and imaging

Wild-type colonies of *Gonium pectorale* (strain CCAC 3275 B, Central Collection of Algal Cultures) were grown axenically in standard *Volvox* medium (SVM) [S1]. Cultures were maintained at 22 ^°^C under approximately 3800 lx illumination on a 14-h/10-h light/dark cycle. Newly released sixteen-cell colonies were selected for recordings, which followed growth toward the next round of cell division. The transition analyzed in the main text occurred while the extracellular matrix (ECM) appeared continuous between cells and the cell number and 4 + 12 arrangement remained unchanged. Later developmental changes in the ECM are described separately in § III.

Micropipettes were pulled from borosilicate glass capillaries (Sutter Instruments; outer diameter 1.0 mm, inner diameter 0.58 mm) using a P-97 puller (Sutter Instruments). The tips were polished using a DMF1000 microforge controller (World Precision Instruments) to an inner diameter of approximately 3 *µ*m. Colonies were transferred to a Petri dish and held individually by gentle aspiration, with suction supplied manually by a 10-mL syringe. The micropipettes were mounted on micromanipulators (Patchstar, Scientifica, UK) carried by a custom rotation stage, allowing face-on or edge-on orientation relative to the optical axis. A layer of mineral oil covered the medium to limit evaporation and the associated convective flows.

The pipette aperture was smaller than a single cell, restricting direct contact to only one peripheral cell. The contacted cell continued to grow and subsequently divided, as did its neighbors. Saddle-shaped colonies were also observed during free swimming. In the continuum model, the localized pipette contact is neglected and the outer boundary is taken to be mechanically free.

Edge-on time-lapse recordings of six colonies were acquired at 1-min intervals on a Zeiss Axiovert 200M inverted microscope with a 40 *×* LD Achroplan objective and a Nikon Z6 II camera. These recordings resolved the evolving colony-body outline used to quantify the flat-to-saddle transition.

A separate set of six colonies was imaged face-on on a Zeiss Axio Observer Z1 inverted microscope with a 40 *×* LD Plan-Neofluar objective and an Axiocam 712 color camera. Bright-field focal series were acquired at ∼5-min intervals over an axial range of approximately 200 *µ*m with a spacing of 2 *µ*m between planes. Each cell was measured in the plane where its boundary was most sharply resolved. These series provided both the projected cell areas used to measure regional growth and the axial positions used in the three-dimensional reconstruction. The corresponding analysis procedures are described in the following subsections.

#### B. Shape analysis and critical size

The colony-body outline was segmented from each edge-on image by intensity thresholding, excluding the micropipette and flagella. Its minimum-area bounding rectangle has long and short sides *L* and *b*, giving the aspect ratio *a* = *b/L*. The pretransition plateau *a*_0_ was estimated from the initial planar portion of each record, and *q* = *a*−*a*_0_ and *ε* = (*L* − *L*_*c*_)*/L*_*c*_ are defined as in Eq. (1) of the main text.

For each colony, *L*_*c*_ and *β* were individually obtained by a least-squares fit of *q* ∝ *ε*^*β*^ for *ε >* 0, with *q* = 0 below onset, to the record before cell division. Across the six colonies, 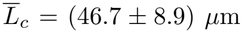 and *β* = 0.52 *±* 0.05 (mean *±* SD). Figure S1 shows the six records with their fits: each colony’s own *L*_*c*_ places its onset at *ε* = 0, while the amplitudes differ between colonies. For the collapse in Fig. 2(b) of the main text, a separate fit of 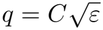 with *L*_*c*_ fixed gives a prefactor *C* for each colony, and *Q* = *q/C*. The amplitude comparison in main text Fig. 3(c) uses *q* itself.

#### C. Cell growth and regional mismatch

The projected cell areas were measured from the face-on focal series described in § I A, each cell in its best-focused plane, and cells were assigned to the central four (*c*) or peripheral twelve (*p*) according to their positions in the fixed arrangement. For each colony, *S*_*α*_(*L*) is the mean cell area of region *α* (*α* = *c, p*) at colony size *L*. The regional linear expansion and relative size *g*_*α*_ and *ξ* are defined by Eq. (2) of the main text, with Δ*g* = *g*_*p*_ − *g*_*c*_. The reference mean areas 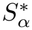 at *L*_*_, the smallest colony size covered by all six records, are obtained by linear interpolation of each record.

**FIG. S1.**
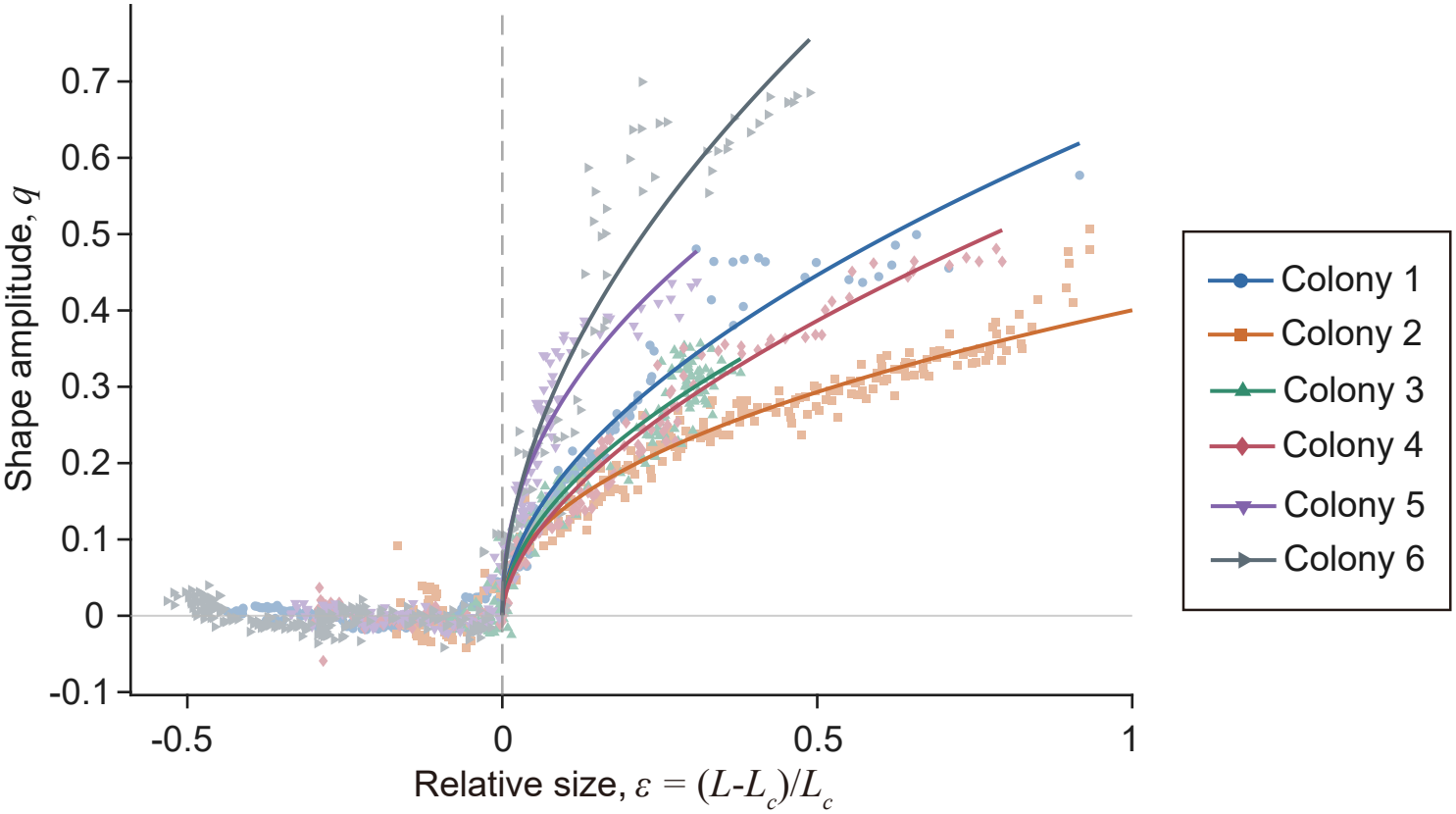
Individual shape records. Shape amplitude *q* versus relative size *ε* for the six edge-on colonies. Lines show the power-law fits for *ε >* 0.

**FIG. S2.**
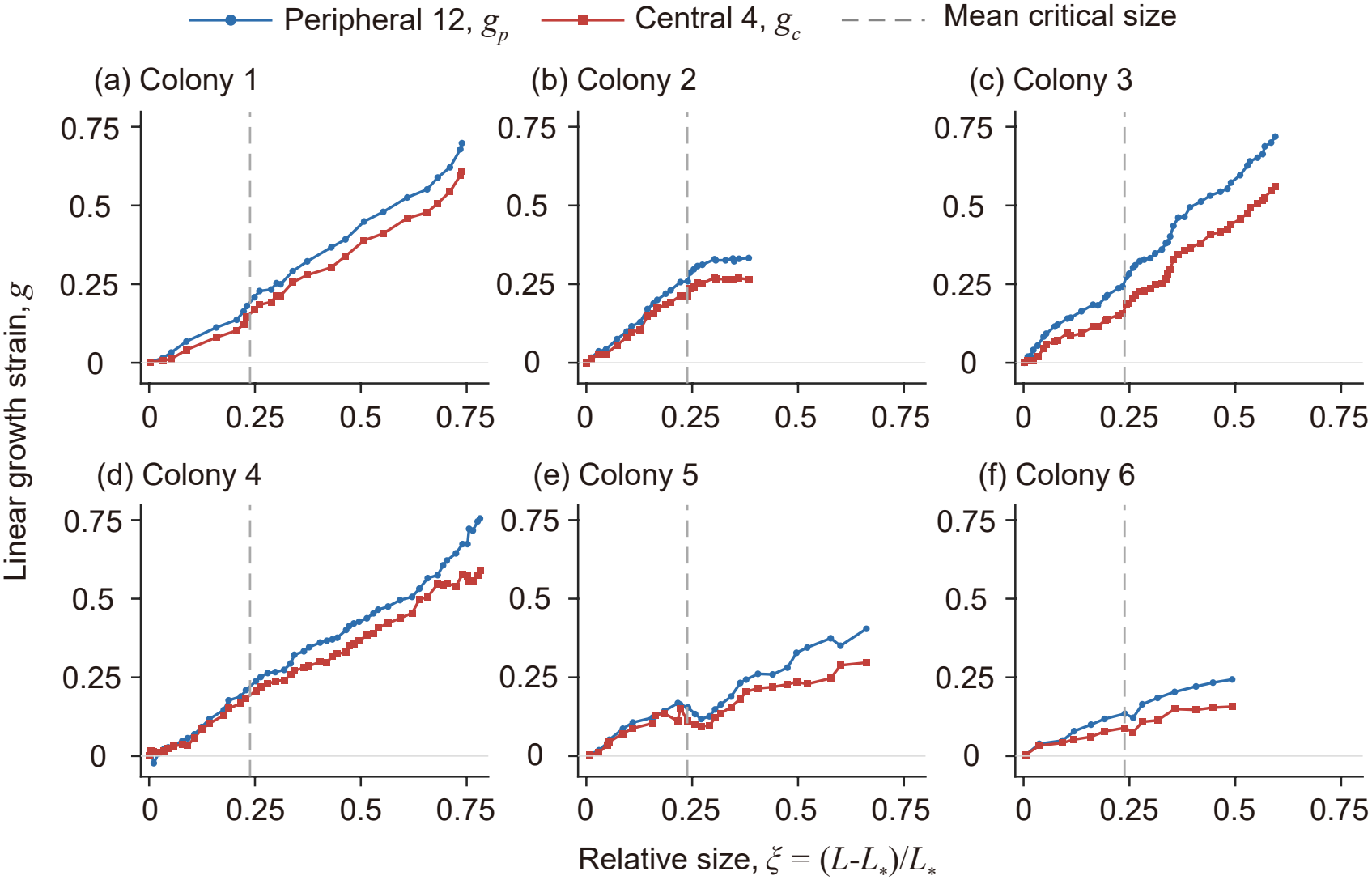
Regional growth in individual colonies. (a)–(f) Cumulative linear expansion of the twelve peripheral (*g*_*p*_, blue) and four central (*g*_*c*_, red) cells in each of the six face-on records, versus *ξ* = (*L − L*_*\**_)*/L*_*\**_. Dashed lines mark 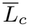. The averages in Fig. 2(c) of the main text use only the range shared by all six colonies.

Figure S2 shows the individual growth curves. For the averages in Fig. 2(c) of the main text, each colony’s curves are interpolated onto a common grid in *ξ*, over the range covered by all six records, and averaged with equal weight; Δ*g* is computed within each colony before averaging, and the bands show the SD across colonies.

#### D. Three-dimensional reconstruction

Cell-center positions were reconstructed from the face-on focal series of § I A. For each cell, its center in the best-focused image gave the lateral coordinates, and the corresponding focal plane gave the axial coordinate. These coordinates were used as centroids of the sixteen cells in a least squares fit to generate a plane for the entire colony. We denote the heights relative to this plane by *ζ*_*j*_ and the azimuths about the colony centroid by *θ*_*j*_.

Each colony was analyzed separately using its twelve peripheral cells. For each *m* = 1, …, 6, a separate least-squares fit at the measured azimuths gave

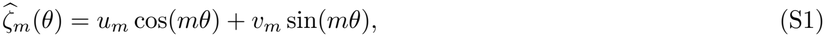

where the hat denotes the fitted height. For *m* = 0, the fitted value was the mean peripheral height. The single-mode explained fraction was defined as

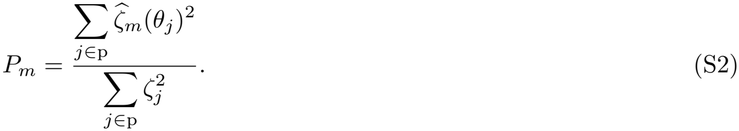

Plane subtraction removes an overall height offset and rigid tilt from all sixteen cells; it does not force *P*_0_ or *P*_1_ of the peripheral subset to vanish. Because the measured azimuths are not uniformly spaced, the separately fitted harmonics are generally not orthogonal, and their explained fractions need not sum to unity.

For each colony, the peripheral *m* = 2 fit was written as *A* cos[2(*θ* − *θ*_0_)], with 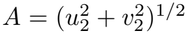. Figure S3 shows the unnormalized heights of all sixteen cells and the individual fits. The central heights were fitted with the same *m* = 2 phase as the periphery, allowing only the amplitude to vary. Central cells therefore enter the plane fit but are excluded from the peripheral harmonic fits.

**FIG. S3.**
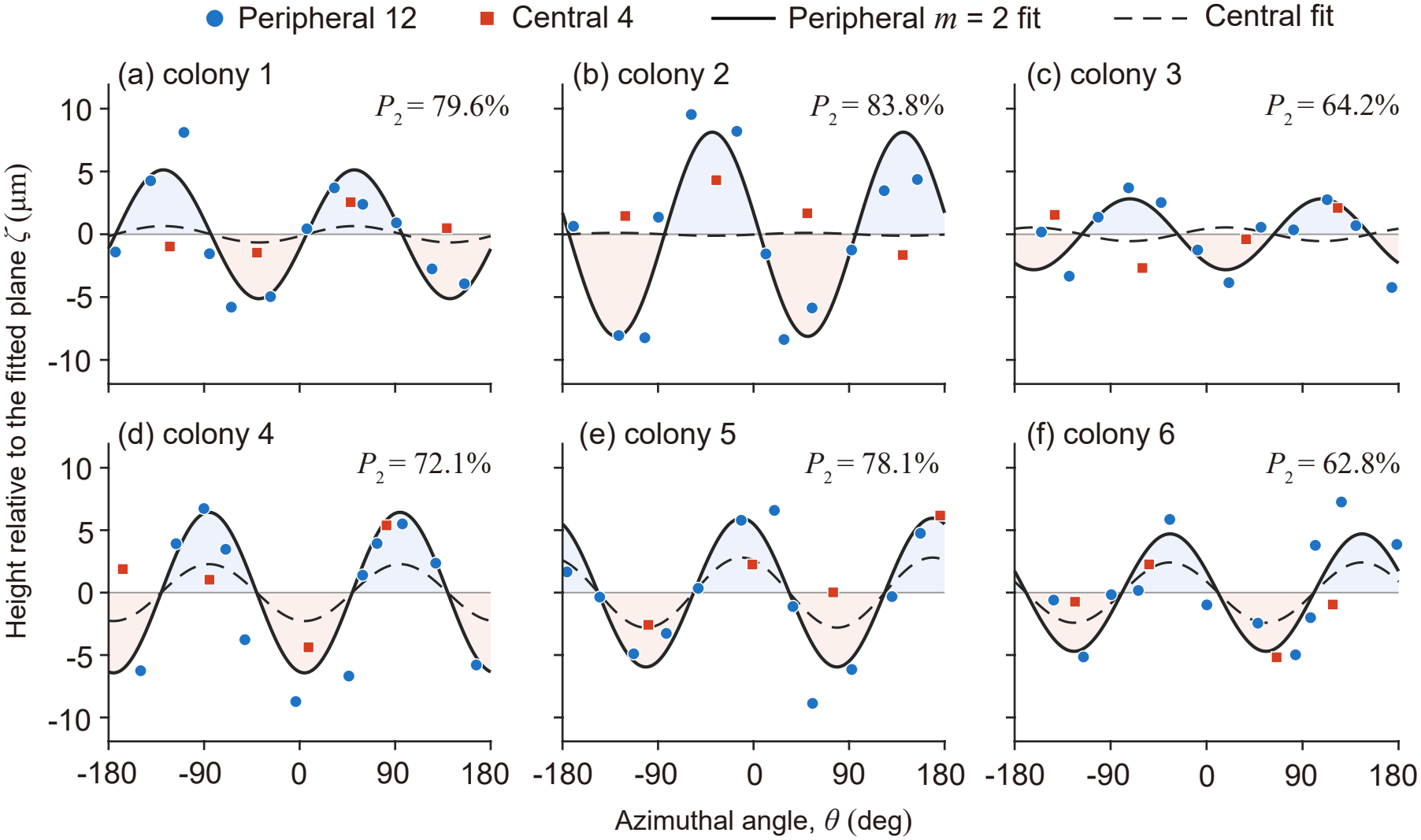
Saddle profiles of six individual colonies. Cell-center heights relative to a plane fitted to all sixteen cells are plotted against azimuth. Blue circles: twelve peripheral cells; red squares: four central cells. Solid curves show the peripheral *m* = 2 fits; dashed curves show central fits with their phase fixed by the peripheral fit.

For Fig. 3(b) of the main text, each colony’s peripheral data set was plotted as *ζ*_*j*_*/A* against *θ*_*j*_ − *θ*_0_. These normalized peripheral data sets were pooled across all colonies and displayed as individual points and in 30^°^ bins. Central cells were not included in this figure.

Figure S4 compares *P*_*m*_ across the six colonies. The *m* = 2 fit accounts for the largest fraction in every colony, with *P*_2_ = (73 *±* 9)% (mean *±* SD across colonies). This establishes the reproducibility of the saddle symmetry using cell-center positions independently of the outline-based amplitude *q*.

**FIG. S4.**
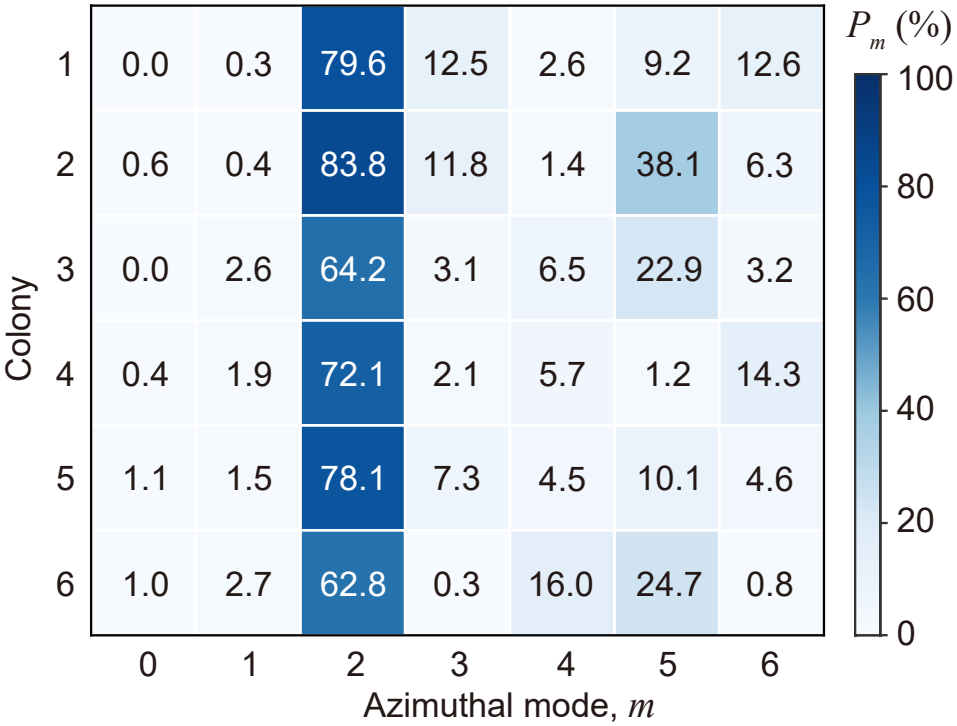
Azimuthal mode content across six colonies. Rows correspond to colonies and columns to separately fitted modes *m* = 0, …, 6. Color and numbers show *P*_*m*_ as a percentage of the squared peripheral height signal explained by each mode after subtraction of the colony-wide best-fit plane.

#### E. Division imaging on the chamber floor

For the division sequence in Fig. 4 of the main text, colonies were immobilized on the floor of a sample chamber rather than held by a micropipette, which is not suited to the late stages of the cycle. Once the 16 daughter colonies have formed [Fig. 1 of the main text], they are linked to one another only through compliant ECM tethers (§ III), which cannot maintain their spatial arrangement; held by a micropipette, the daughters drift out of the focal plane and the field of view. Colonies were instead immobilized by light-induced flagellar adhesion, as reported for *Chlamydomonas* [S2]. The chamber was built from a glass slide and a coverslip separated by double-sided tape, giving a depth of ∼2 mm. Under illumination, the flagella of colonies settling on the glass adhere to it, holding each colony in place for long-term recording. Adhesion keeps the colony flat against the substrate, so these recordings are used only to follow the arrangement of the daughter colonies, not the growth-driven saddle described in the main text. Colonies remained within the field of view during division, allowing daughter formation and reorientation to be followed continuously. Bright-field time-lapse recordings were acquired at 1-min intervals with the microscope, objective, and camera described in § I A.

### II. ELASTIC MODEL

#### A. Incremental equations

We use the geometry, growth field and effective coefficients of the main text. Growth is slow compared with elastic relaxation, so the sheet is in mechanical equilibrium for the current *g*; growth is uniform across the thickness and imposes no preferred curvature. With lengths in units of *R, w* = *ζ/R*, Φ = *χ/*(*KR*^2^) and Γ = *D/*(*KR*^2^) as in the main text, the growth-modified Föppl–von Kármán equations [S3] for the total stress function are

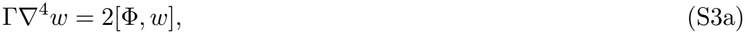

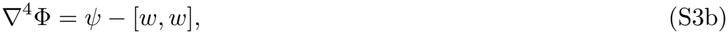

where [*f, g*] = (*f*_,*xx*_*g*_,*yy*_ + *f*_,*yy*_*g*_,*xx*_ −2*f*_,*xy*_*g*_,*xy*_)*/*2, so that [*w, w*] is the (scaled) Gaussian curvature of the sheet, and *ψ* is the Gaussian curvature prescribed by growth. Because uniform growth merely rescales each region, *ψ* vanishes within the center and the periphery and is localized at the interface *ρ* = *λ*.

The flat state *w* = 0 satisfies the first of Eqs. (S3) identically, while the second reduces to ∇^4^Φ^(0)^ = *ψ*. Rather than solving this equation, the End Matter obtains the corresponding stresses 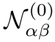 directly from the displacement. The Airy representation, *N*_*xx*_ = Φ_,*yy*_, *N*_*yy*_ = Φ_,*xx*_ and *N*_*xy*_ = −Φ_,*xy*_, gives

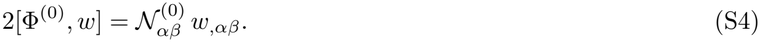

We compare the buckled sheet with a flat reference state at the same prescribed growth. Writing 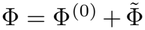 and subtracting ∇^4^Φ^(0)^ = *ψ* from the second of Eqs. (S3) cancels *ψ*. Using Eq. (S4) in the first equation then gives the incremental equations of the main text:

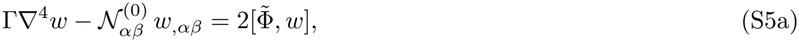

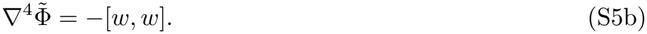

The growth source *ψ* cancels from the incremental compatibility equation, while differential growth remains through the prestress 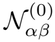.

#### B. Mode selection

We solved the radial mode equation with the interface and free-edge conditions of the End Matter for *m* = 0, …, 10 at *λ* = *ν* = 1*/*2. At the center, smoothness requires 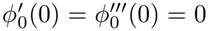 for *m* = 0 and 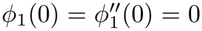 for *m* = 1; the conditions for *m* ≥ 2 are given in the End Matter.

After excluding rigid translation (*m* = 0, constant height) and rigid tilt (*m* = 1, *w* ∝ *ρ* cos *θ*, a tilted plane), no positive buckling eigenvalue was found for *m* = 0 or 1. Over 2 ≤ *m* ≤ 10, the thresholds in Table S1 increase monotonically with *m*. This trend is consistent with the leading *m*^4^ angular bending term, compared with the *m*^2^ hoop-compression term. The lowest threshold, *η*_*c*_(2) = 9.16, corresponds to the observed saddle. Increasing the numerical resolution did not change the reported digits.

**TABLE S1.** Smallest positive eigenvalue *η*_*c*_(*m*) for *λ* = *ν* = 1*/*2. Dashes indicate that none was found after removing rigid translation and tilt.

| *m* | 0 | 1 | 2 | 3 | 4 | 5 | 6 | 7 | 8 | 9 | 10 |
|---|---|---|---|---|---|---|---|---|---|---|---|
| *η*<sub>*c*</sub>(*m*) | — | — | 9.16 | 19.4 | 33.7 | 52.1 | 74.6 | 101 | 132 | 166 | 205 |

#### C. Amplitude prediction and comparison

The critical profile reaches its largest magnitude at the rim, where *ϕ*_*c*_(1) = 1. The maximum and minimum midsurface heights are thus *R*|*A*| and −*R*|*A*| to leading order, giving a total height range 2*R*|*A*|. Projection along any direction in the original plane of the colony retains this range, irrespective of the azimuthal viewing direction.

For the outline measurement, we retain the measured planar baseline *a*_0_ and take the in-plane span to be *L* = 2*R* to leading order. The short side of the enclosing rectangle is then

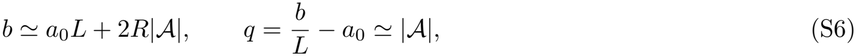

and the amplitude equation gives, above onset,

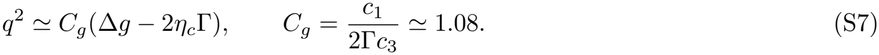

The amplitude comparison assumes Γ ≃ Γ_*c*_ over the near-onset window. Growth and shape were recorded in separate cohorts, so the mean critical size 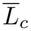 from the side-view measurements is used as the onset input for all six face-on growth records. For each fitted mismatch curve Δ*g*_*i*_(*L*), we evaluate its critical mismatch 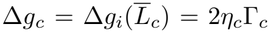 and calculate the amplitude from Eq. (S7). The six predicted amplitudes are then averaged to give the prediction shown in Fig. 3(c) of the main text. The geometric conversion introduces no additional fitted prefactor.

### III. ECM REMODELLING IN LATER STAGES

Figure S5 shows ECM morphology at different stages of the developmental cycle summarized in Fig. 1 of the main text. The panels show different colonies rather than successive frames of one colony. In an early colony, Alcian Blue staining reveals a common envelope surrounding the cells [Fig. S5(a)]. Later in growth, a mesh-like intercellular matrix is visible between the cell bodies, including the four central cells [Fig. S5(b)]. A colony containing cells that have begun division shows both this mesh-like pattern and elongated intercellular connections [Fig. S5(c)]. At a more advanced stage, developing daughter colonies remain linked by slender, tether-like structures [Fig. S5(d)]. During subsequent development, peripheral daughter colonies turn out of plane, giving the arrangement shown in Fig. 4 of the main text.

**FIG. S5.**
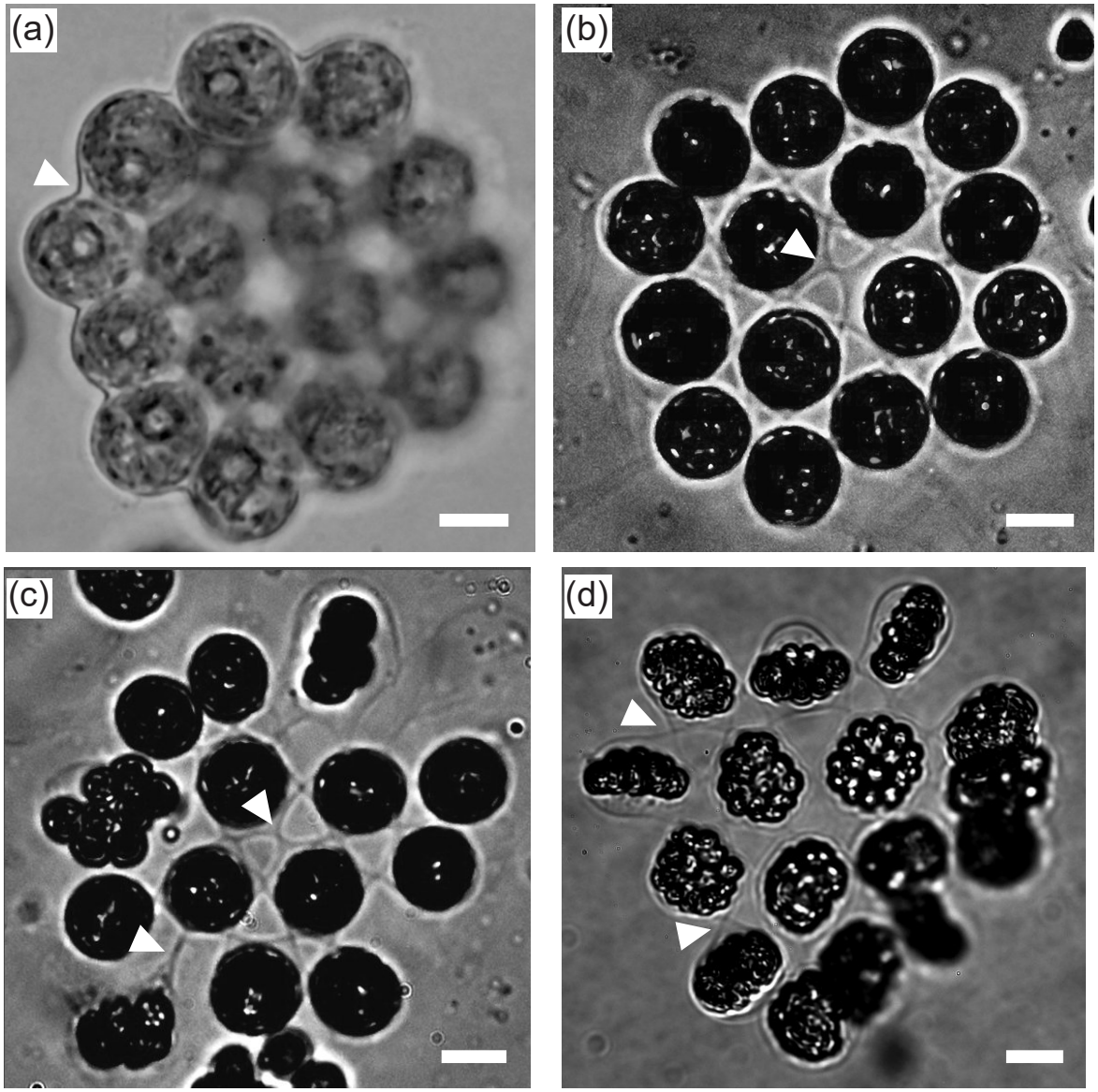
Representative ECM morphologies during *Gonium pectorale* development. (a) Colony imaged after Alcian Blue staining; the arrowhead marks the common outer envelope. (b) Mesh-like intercellular matrix, visible between the central cells (arrowhead). (c) Colony containing dividing cells, showing a mesh-like matrix (upper arrowhead) and an elongated connection (lower arrowhead). (d) Developing daughter colonies linked by tether-like structures (arrowheads). The panels show different colonies and do not form a time-lapse sequence. Brightness and contrast were adjusted to improve matrix visibility; images are displayed in grayscale. Scale bars: 10 *µ*m in (a)–(c) and 20 *µ*m in (d).

These observations motivate a structural interpretation of the reduced effective bending-to-stretching ratio Γ inferred in the main text. The saddle transition analyzed quantitatively occurs during the earlier growth stage, when the matrix appears continuous. At this stage, however, the matrix may already vary in thickness, with thinner regions between neighboring cells. These thin connections could bend more easily while still transmitting in-plane forces, reducing the colony’s resistance to bending relative to stretching. The later mesh and tethers provide morphological support for this possibility.

### IV. SUPPLEMENTAL MOVIE CAPTIONS

**Supplementary Movie 1**: Flat-to-saddle transition viewed from the side. Edge-on time-lapse recording of a sixteen-cell *Gonium pectorale* colony held by gentle micropipette aspiration. The colony grows continuously while retaining its 4+12 cell arrangement and, without further cell division, develops from an initially nearly planar configuration into a pronounced out-of-plane shape over approximately 29 h.

**Supplementary Movie 2**: Axial bright-field scan through a saddle-shaped colony. The focal plane is stepped through the colony at 2-*µ*m intervals, so that cells at different axial positions come successively into focus, illustrating the measurements underlying the three-dimensional reconstruction.

**Supplementary Movie 3**: Division of a colony on the chamber floor, corresponding to Fig. 4(b)–(e) of the main text. The colony is immobilized by light-induced flagellar adhesion and remains flat (§ I E). Each of the sixteen cells forms a sixteen-cell daughter colony; the twelve peripheral daughters come to stand edge-on in a ring around the four central ones. Bright-field time-lapse recorded at 1-min intervals. Scale bar, 10 *µ*m.

## References

[1] J. Huxley, Problems of Relative Growth (The Dial Press, New York, 1932).

[2] D. W. Thompson, On Growth and Form (Cambridge University Press, Cambridge, 1917).

[3] E. K. Rodriguez, A. Hoger, and A. D. McCulloch, Stress-dependent finite growth in soft elastic tissues, J. Biomech. 27, 455 (1994).

[4] E. Efrati, E. Sharon, and R. Kupferman, Elastic theory of unconstrained non-Euclidean plates, J. Mech. Phys. Sol. 57, 762 (2009).

[5] A. Goriely and M. Ben Amar, Differential growth and instability in elastic shells, Phys. Rev. Lett. 94, 198103 (2005).

[6] J. Dervaux and M. Ben Amar, Morphogenesis of growing soft tissues, Phys. Rev. Lett. 101, 068101 (2008).

[7] J. Dervaux, P. Ciarletta, and M. B. Amar, Morphogenesis of thin hyperelastic plates: a constitutive theory of biological growth in the Föppl–von Kármán limit, J. Mech. Phys. Sol. 57, 458 (2009).

[8] Y. Klein, E. Efrati, and E. Sharon, Shaping of elastic sheets by prescription of non-euclidean metrics, Science 315, 1116 (2007).

[9] J. Kim, J. A. Hanna, M. Byun, C. D. Santangelo, and R. C. Hayward, Designing responsive buckled surfaces by halftone gel lithography, Science 335, 1201 (2012).

[10] T. Mora and A. Boudaoud, Buckling of swelling gels, Eur. Phys. J. E 20, 119 (2006).

[11] H. Liang and L. Mahadevan, The shape of a long leaf, Proc. Natl. Acad. Sci. USA 106, 22049 (2009).

[12] H. Liang and L. Mahadevan, Growth, geometry, and mechanics of a blooming lily, Proc. Natl. Acad. Sci. USA 108, 5516 (2011).

[13] T. Savin, N. A. Kurpios, A. E. Shyer, P. Florescu, H. Liang, L. Mahadevan, and C. J. Tabin, On the growth and form of the gut, Nature 476, 57 (2011).

[14] U. Nath, B. C. Crawford, R. Carpenter, and E. Coen, Genetic control of surface curvature, Science 299, 1404 (2003).

[15] E. Coen, A.-G. Rolland-Lagan, M. Matthews, J. A. Bangham, and P. Prusinkiewicz, The genetics of geometry, Proc. Natl. Acad. Sci. USA 101, 4728 (2004).

[16] Y. Zhang, O. Y. Cohen, M. Moshe, and E. Sharon, Geometrically frustrated rose petals, Science 388, 520 (2025).

[17] S. Höhn, A. R. Honerkamp-Smith, P. A. Haas, P. K. Trong, and R. E. Goldstein, Dynamics of a Volvox embryo turning itself inside out, Phys. Rev. Lett. 114, 178101 (2015).

[18] E. R. Hanschen, T. N. Marriage, P. J. Ferris, T. Hamaji, A. Toyoda, A. Fujiyama, R. Neme, H. Noguchi, Y. Minakuchi, M. Suzuki, et al., The Gonium pectorale genome demonstrates co-option of cell cycle regulation during the evolution of multicellularity, Nat. Comm. 7, 11370 (2016).

[19] M. D. Herron and R. E. Michod, Evolution of complexity in the volvocine algae: transitions in individuality through Darwin’s eye, Evolution 62, 436 (2008).

[20] H. De Maleprade, F. Moisy, T. Ishikawa, and R. E. Goldstein, Motility and phototaxis of Gonium, the simplest differentiated colonial alga, Phys. Rev. E 101, 022416 (2020).

[21] R. A. Harper, The structure and development of the colony in Gonium, Trans. Am. Micro. Soc. 31, 65 (1912).

[22] D. L. Kirk and M. M. Kirk, Protein synthetic patterns during the asexual life cycle of Volvox carteri, Dev. Biol. 96, 493 (1983).

[23] See Supplemental Material at http://link.aps.org/supplemental/10.1103/XXX for further experimental videos and details of the theoretical analysis.

[24] L. D. Landau, L. Pitaevskii, A. M. Kosevich, and E. M. Lifshitz, Theory of elasticity, Vol. 7 (Elsevier, 2012).

[25] E. Memet, F. Hilitski, M. A. Morris, W. J. Schwenger, Z. Dogic, and L. Mahadevan, Microtubules soften due to cross-sectional flattening, eLife 7, e34695 (2018).

[26] Y. Cohen, A. Pandey, Y. Zhang, C. Maor, and M. Moshe, Curvature potential formulation for thin elastic sheets, Phys. Rev. Lett. 137, 188201 (2026).

[27] T. Kubo, S. Kaida, J. Abe, T. Saito, H. Fukuzawa, and Y. Matsuda, The Chlamydomonas hatching enzyme, sporangin, is expressed in specific phases of the cell cycle and is localized to the flagella of daughter cells within the sporangial cell wall, Plant Cell Physiol. 50, 572 (2009).

[28] N. Ueki and I. Nishii, Controlled enlargement of the glycoprotein vesicle surrounding a Volvox embryo requires the invb nucleotide-sugar transporter and is required for normal morphogenesis, Plant Cell 21, 1166 (2009).

[29] Z. Wang, S. K. Birwa, J. Y. Y. Chui, K. C. Leptos, and R. E. Goldstein, Data for “Saddle Buckling of a Multicellular Alga Driven by Differential Growth” (2026), zenodo, doi:10.5281/zenodo.24037401.

## References

[S1] D. L. Kirk and M. M. Kirk, Developmental biology 96, 493 (1983).

[S2] C. T. Kreis, M. Le Blay, C. Linne, M. M. Makowski, and O. Bäumchen, Nature Physics 14, 45 (2018).

[S3] J. Dervaux and M. Ben Amar, Physical review letters 101, 068101 (2008).

